# Cryo-EM structure of the *Rhodobaca bogoriensis* RC-LH1-PufX dimeric complex at 2.9 Å

**DOI:** 10.1101/2022.02.25.481955

**Authors:** Dmitry A. Semchonok, Marina I. Siponen, Christian Tüting, Quentin Charras, Fotis L. Kyrilis, Farzad Hamdi, Yashar Sadian, Colette Jungas, Panagiotis L. Kastritis

## Abstract

The reaction centre-light harvesting 1 (RC-LH1) complex is essential for converting light into proton motive force in photosynthetic bacteria. RC-LH1 is a monomer in most purple bacteria, but in Rhodobacter species, it is a dimer. Its assembly depends on an accessory polypeptide (PufX) and, ultimately, on photosynthetic growth. To date, knowledge on the RC-LH1-PufX structure, where the dimer has two incomplete ‘C’-shaped antenna rings surrounding an RC, is mainly limited to the model organism *Rhodobacter sphaeroides*. Here we present a cryo-electron microscopy structure at 2.9 Å from *Rhodobaca bogorensis* strain LBB1. RCs are surrounded by 30 antennas and incorporate protein Y and PufX. RCs are stably connected by PufX, which self-interacts, electrostatically attracts cytochrome *c2* (cyt *c2*) and forms extensive networks with co-factors. This structure underlines coordinated energy transfer in a combinatorial manner, providing a basis to describe bacterial photosynthesis within a dimeric photosynthetic apparatus.

## Introduction

In photosynthetic bacteria, the early steps of photosynthesis (light harvesting and its subsequent conversion into a stable charge separation), and the late ones (cyclic electron transport and accumulation of an electrochemical gradient of protons), depend upon the relationship between the reaction centre (RC) and the *bc*1 complex^1, 2, 3, 4^. The light captured by the RC is used to reduce a membrane-embedded quinone molecule, which then diffuses to the cytochrome *bc*1 complex to be oxidised. For the capture of the solar energy, the photosynthetic purple bacteria use two types of light-harvesting complexes, LH1 antenna, close to the reaction centre and present in a fixed ratio with the RC (depending on the species^5, 6, 7, 8^) and the peripheral LH2 complexes, which are subjected to regulation regarding oxygen and light concentration.

The supramolecular architecture of the RC-LH1 supercomplex shows a ring of LH1 antennae around the reaction centre of 14, 15 or 16 αβ-LH1 subunits depending on whether the LH1 antenna ring is open or closed^9,10,11^. Fifteen αβ-LH1 subunits are found in *Rhodospeudomonas palustris* where the ring is only partially closed^12^. Few high-resolution structures have been described for *Thermochromatium tepidum*^13, 14^, *Blastochloris viridis*^15^ and *Rhodospirillum rubrum*^16^, where the LH1 antenna of 16 αβ-subunits exhibits a closed ring around the RC. In Rhodobacter species, RC-LH1 dimeric supercomplexes have been reported, where two incomplete ‘C’-shaped (S-shaped array) antenna rings of 14 LH1s each surround an RC, confirming the dual role played by PufX as a linker and causing interruption of the ring of LH^17, 18, 19, 20, 21^. The structural role of PufX in *R. sphaeroides* was recently described by solving cryo-EM structures of RC-LH1 monomer and dimer complexes of a modified dark-grown strain1^22, 23^.

Here, we characterise the RC-LH1-PufX dimeric supercomplex, the only pigment-protein assembly present in Rhodobaca (*Rca*.*) bogoriensis* at 2.9 Å, resolved by cryo-electron microscopy (cryo-EM). The model informs on the non-covalent interactions among 70 component polypeptide chains and 154 co-factor molecules, localising 45174 atoms in total. The RCs are surrounded by 30 antennas that bind 58 spheroidenones (Spno) and 60 bacteriochlorophyll molecules (BChl a). Each RC incorporates protein Y, 4 BChl a, 3 pheophytins (Pheo), 5 quinones, a nonheme-iron atom, a Spno and 4 phospholipids. RCs are connected by PufX, which self-interacts, electrostatically attracts electron donors/acceptors, binds chlorophylls and lipids, substitutes Spno, and, ultimately, links the RC-LH1 complexes. This structure underlines coordinated energy transfer in a combinatorial manner, and overall, provides a basis to describe bacterial photosynthesis within a dimeric photosynthetic apparatus.

## Results and Discussion

### Visualisation of the RC-LH1-PufX dimer in native *Rca. bogoriensis* photosynthetic membranes

Electron microscopy (EM) observations of *Rca. bogoriensis* cells highlight a unique organization of the lipid membranes **(Fig. 1a-b)** as the bacteria appear to display intracytoplasmic membranes, *i*.*e*., chromatophores, organized alongside the cell wall inner membrane. Freeze-fracture EM shows that RC-LH1-PufX dimers are embedded in chromatophores **(Fig. 1c-d)**. *Rca. bogoriensis c*hromatophores appear near-spherical, an exemplary architecture for maximizing the probability to capture photons while remaining in a confined surface, therefore, extending the total photosynthetic surface area. Freeze fracture also highlights the peculiar organisation of the RC-LH1-PufX dimers in both intracytoplasmic and cytoplasmic membrane (**Fig. 1c** and **Fig. 1d**). Indeed, the dimers appear to form an ordered organisation. Such a macromolecular organisation may be a key to optimising electron transfer by creating quinone “highways’’ between the RC-LH1-PufX and *bc*1 complexes. Indeed, the reduced quinone released at the QB site of the reaction centre must be rapidly replaced by an oxidised quinone in order not to block the cyclic electron transfer since under anaerobic growth conditions, one oxidised quinone is present for every 50 reduced quinones^24^.

**Figure 1.**
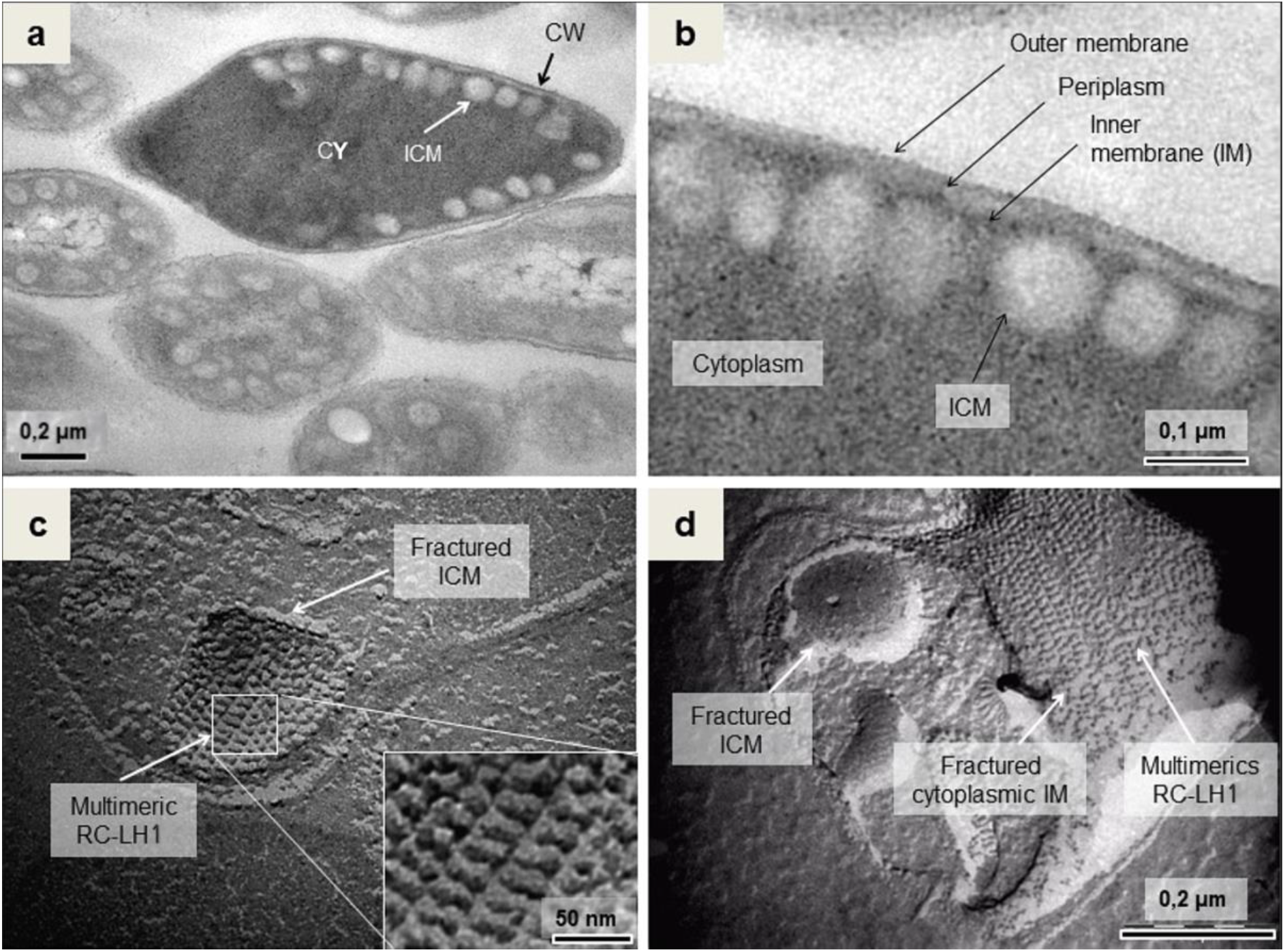
Cellular and photosynthetic membrane macromolecular organisation in *Rca. bogorensis* strain LBB1. (a) Negative-staining EM observation of resin embedded *Rca. bogoriensis* bacteria cells. (CY) Cytoplasm, (ICM) Intracytoplasmic membrane, (CW) cell wall. (b) Close view of the *Rca. bogoriensis* cell (from (a)) highlighting the cell wall organisation and the intracytoplasmic membranes (ICM). (c) Freeze-fracture EM view of the *in vivo* ICM, inside which, the RC-LH1-PufX dimers are inserted. The close view of the membrane displays the well-ordered organisation of the RC-LH1-PufX dimers. (d) Freeze-fracture EM view of ICM together with the cytoplasmic side of the inner membrane, inside which the RC-LH1-PufX dimers are also inserted.

### Cryo-EM structure of the native *Rca. bogoriensis* RC-LH1-PufX complex

After defining the *in vivo* relevance of the RC-LH1-PufX dimer, we optimised the isolation procedure utilising sucrose gradient purification (**Fig. 2a**) with dodecyl-β-D-maltoside (DDM) solubilisation of the *Rca. bogoriensis* photosynthetic membranes. The sucrose gradient shows a stable dimer compared to those retrieved from other purple bacteria **(Fig. 2a)**. After vitrification and optimising the cryo-conditions, we set up a special acquisition protocol (see Methods) to visualise both top and side views, amounting to 120 e-/Å^2^ per complete movie **(Fig. 2b)**. This allowed particle alignments by increasing the signal-to-noise ratio and taking advantage of dose weighting, 2D class averages showed high-resolution features in unrestrained projections **(Fig. 2c)**.

**Figure 2.**
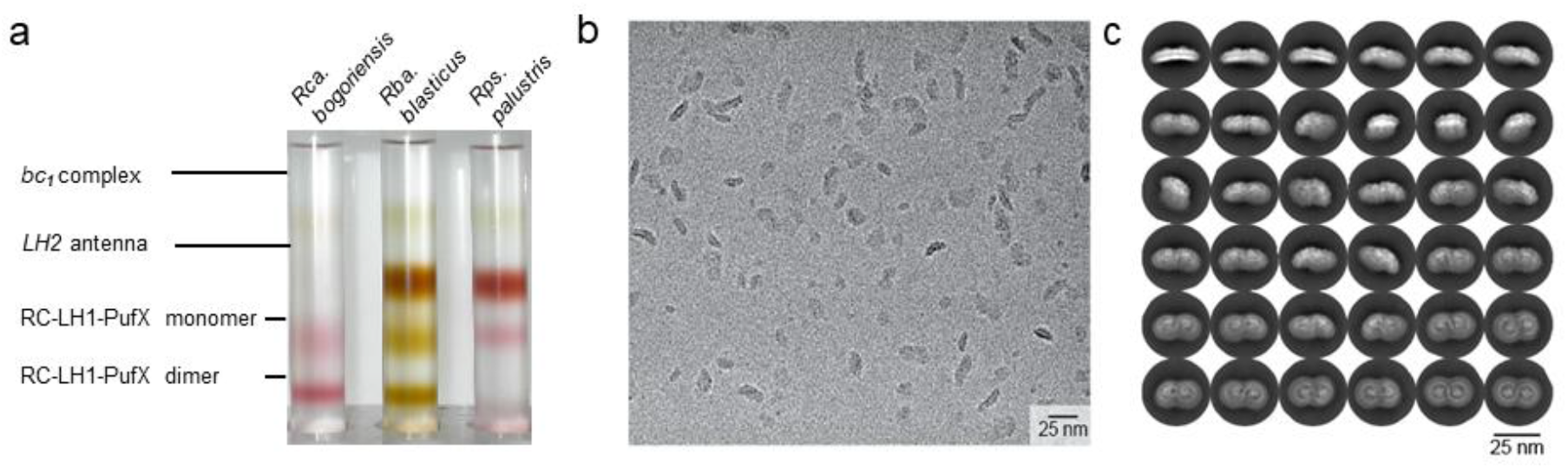
Successful dimer purification, vitrification and unrestrained 2D classification of the RC-LH1-PufX dimer. (a) Comparative sucrose gradient from solubilised chromatophores from 3 different purple bacteria. (b) cryo-electron micrograph derived from summing 120 frames, each exposed for 1 e-/Å^2^/sec, showing a variety of orientations for RC-LH1-PufX dimer in vitreous ice. (c) 2D class averages exhibit a plethora of 2D projections of the RC-LH1-PufX dimer.

We retrieved a cryo-EM map at 2.9 Å resolution (FSC=0.143) with a B-factor of −99.6 Å^2^ for the endogenous, wild-type RC-LH1-PufX dimer from *Rca. bogoriensis* after image processing (**Fig. 3a, Extended Data Fig. 1, Extended Data Fig. 2a-b**). Additional asymmetric reconstructions resolved a cryo-EM map at 3.0 Å (FSC=0.143) (**Extended Data Fig. 2c-d)**, virtually identical to its C2 symmetric counterpart (**Extended Data Fig. 2a-b**), revealing a naturally occurring symmetry of the *Rca. bogoriensis* RC-LH1-PufX dimer. The reconstructed map quality was high, as revealed by the extensive coverage of distinct views (**Extended Data Fig. 3a**), while the mask was covering lower resolution voxels, *e*.*g*., corresponding to the detergent (**Extended Data Fig. 3b-c)**. The atomic structure has a diameter of 220.5 Å, a surface area of 0.3245E+05 Å^2^, and an overall volume of 0.7960E+06 Å^3^, while the association of the RC-LH1 monomers results in a concave surface for the supercomplex with an angle of 166^0^ **(Fig. 3b)**. Our reported RC-LH1-PufX dimer is, at the time of publication, the only complex from a wild-type species observed in photosynthetic membranes with freeze-fracture electron microscopy **(Fig. 1)**. The RC-LH1-PufX dimeric surface is substantially flatter than those previously reported for an LH1-deficient strain of *Rhodobacter sphaeroides* grown in the dark^23^. Although flexibility can rationalise this discrepancy^25^, 2D class averages showed a remarkably uniform signal (**Fig. 2c, Extended Data Fig. 2e)** and local resolution analysis indicates that variation in the endogenous RC-LH1 dimer is confined at the supercomplex interface periphery (**Extended Data Fig. 2f**). The substantially less bent configuration of the *Rca. bogoriensis* RC-LH1 dimer can have various biological consequences, including, but not limited to, regulating the size, shape, and curvature of the wild-type spherical chromatophores previously observed^26^ **(Fig. 1)**, phase-separating membrane-embedded photosynthetic complexes^27^, assisting the formation of their observed lattice^9^, and regulating possible effects on the excitation transfer, previously observed in other species^28^. Finally, due to the curvature, the dimer has a larger surface and spatial angle coverage and is possibly critical at increasing depths, capturing more scattered photons as light in water comes from different directions. The obtained resolution allowed to localize 70 protein components, 154 pigment molecules and to model the positions of 45174 atoms (**Extended Data Table 1**). Map quality allows clear visualising the special pair of chlorophylls in the RC and their local environment **(Fig. 3c)** and resolving various co-factors within the structure **(Fig. 3d)**.

**Figure 3.**
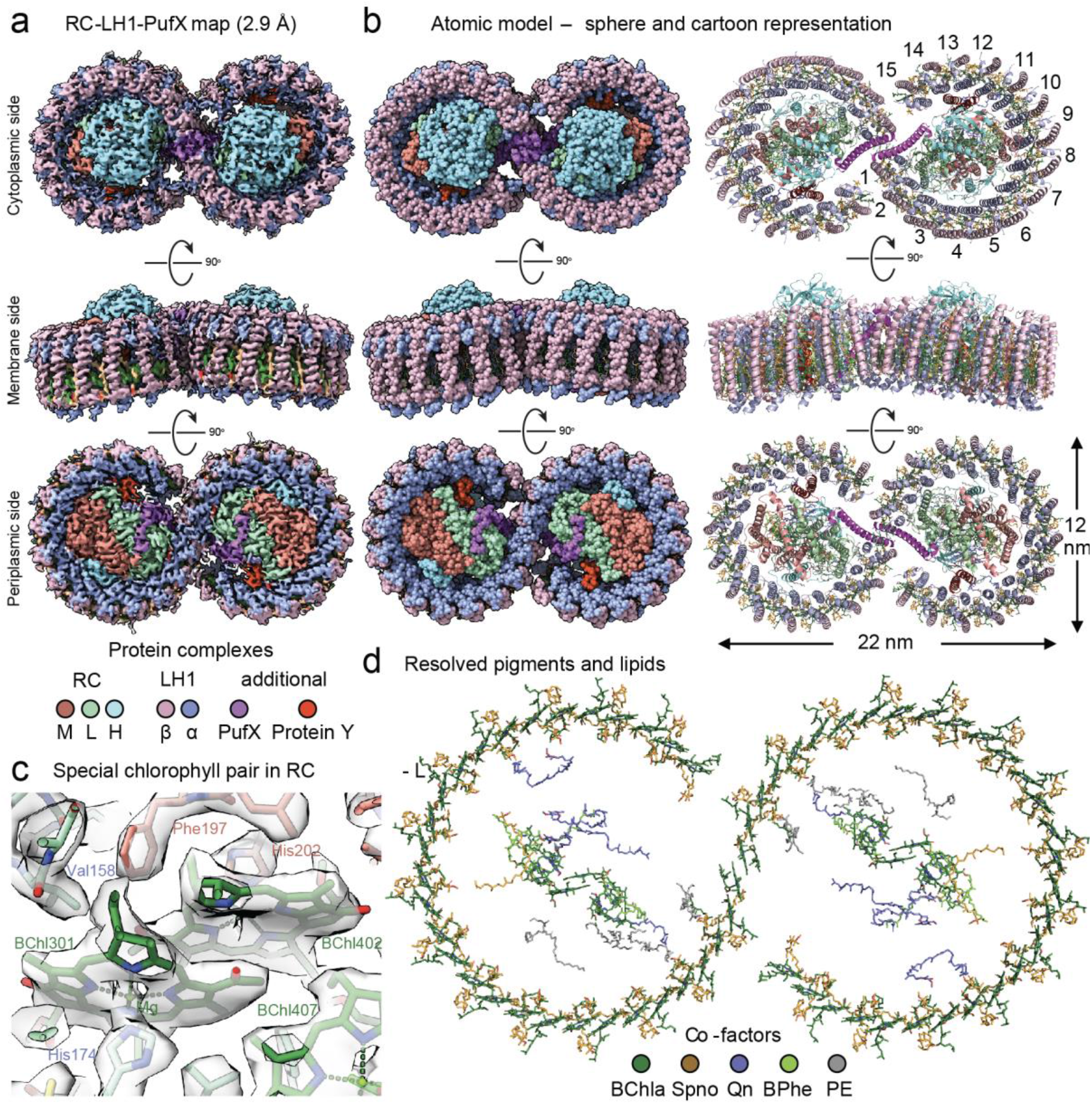
Overview of the *Rca. bogoriensis* RC-LH1-PufX dimer. (a) cryo-EM map with coloured protein components; (b) Sphere and cartoon representation of the atomic model satisfying the derived cryo-EM density; (c) Resolution of the cryo-EM map, focusing on the chlorophyll pair within the RC and the proximal side-chains; (d) top view of the modelled co-factors within the RC-LH1-PufX dimer.

### The unique architecture of LH1

Fifteen LH1 subunits curve around each RC (**Fig. 3b**) in an arrangement reminiscent of the mathematical symbol of infinity (∞). The 15^th^ LH1 subunit replaces the Protein Z previously reported for *R. sphaeroides*^23^, signifying the presence of the two additional LH1 subunits, resulting in a higher antenna density surrounding the *Rca. bogoriensis* RCs. This high antenna density directly impacts excitation transfer mechanisms within the RC-LH1 dimer. Surface accessibility calculations show that the additional LH1 subunits almost occlude the dimeric interface next to the quinone channel. This interaction surface is formed by the 15^th^ LH1 subunit and the 1^st^ and 2^nd^ LH1 subunits from the symmetric RC-LH1 monomer. Partial occlusion of the dimeric interface can well be a mechanism regulating the quinone/quinol exchange, varying the accessibility of the formed channel, as recently observed in *Rh. Palustris* RC-LH1 monomer^7^. 14 out of 15 LH1 subunits bind 4 pigment molecules, *i*.*e*., 2 bacteriochlorophylls a (BChl a) and 2 spheroidenones (Spn or Spno), the latter never previously resolved in a photosystem structure. The number of carotenoids matches those to *R. sphaeroides*, where Spno is replaced by methoxy-neurosporene^23^.

The expected interactions of α and β LH1 chains with each BChl a are derived (reviewed in^29^) and include the ligation of Mg of α and β BChls by His (α-His32 and β-His39, respectively) and the hydrogen bonding of their C3 acetyl carbonyls with Trp (α-Trp42 and β-Trp48, respectively) **(Fig. 4a-b)**. Interestingly, α-Trp45 is in proximity with β-BChl a (dmin=3.6 Å) and not α-BChl a (dmin=6.0 Å), stabilising the α/β subunits. Spnos are asymmetrically distributed across the subunit, with one passing through the heterodimer utilising the hydrophobic tail. At the same time, the second Spno localises on the outer surface of the LH1 in an almost parallel fashion (**Fig. 4b)**.

**Figure 4.**
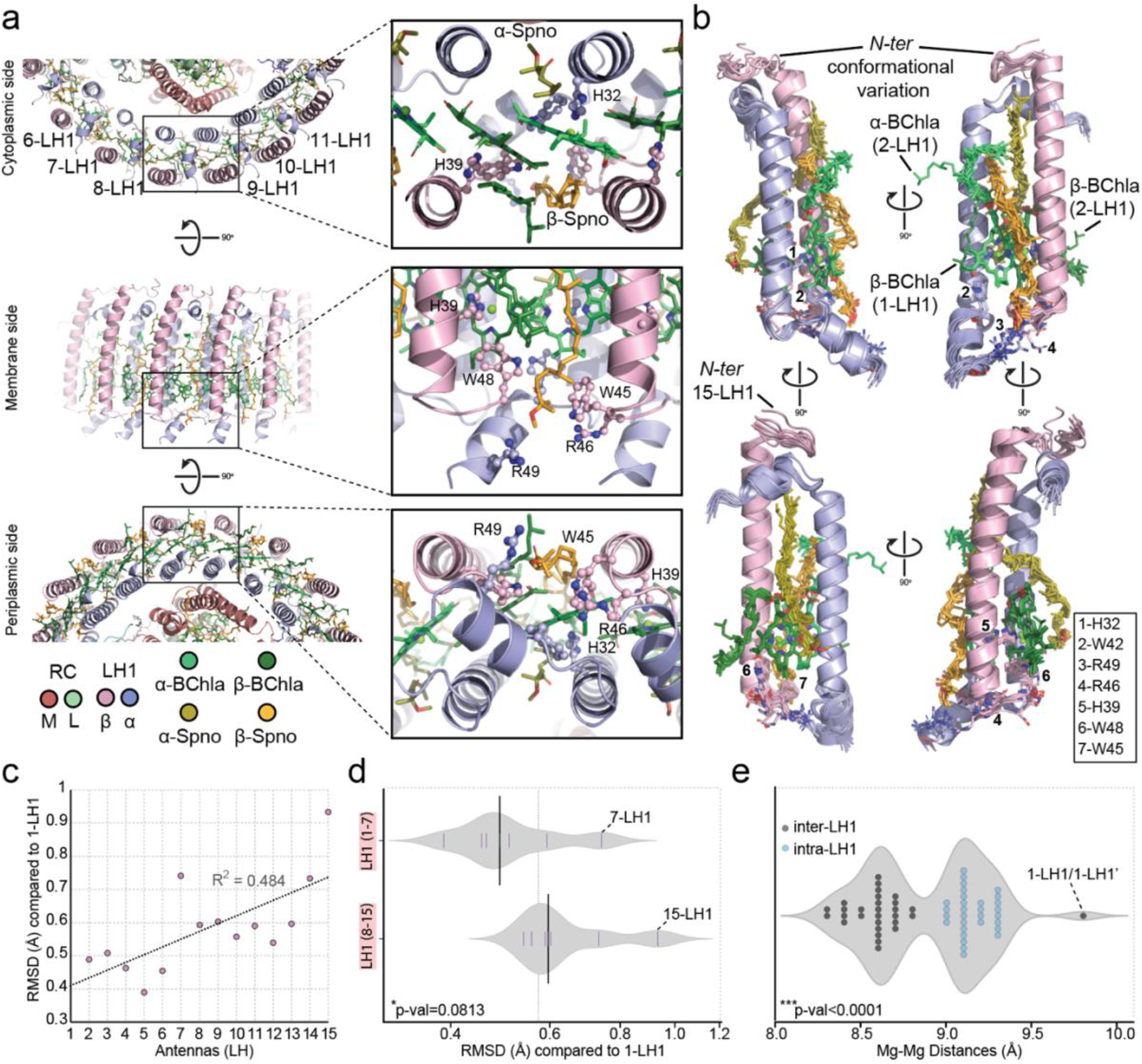
The unique features of the LH1 of *Rca. bogoriensis* dimer. (a) Pigment - protein organisation within LH1 subunits; (b) Superposition of all 15 LH1 subunits showing the highly conserved secondary structure and binding of pigments; However, 1-LH1 and 2-LH1 exhibit conformational changes in the BChl a tails, and the *N-ter* of 15-LH1 shows substantial displacement; (c) Increased conformational deviation from 1-LH1 as a function of LH1 subunit number; (d) Violin plot describing significant differences in conformational similarity of LH1 subunit groups with 1-LH1; (e) Violin plot describing the distribution of Mg-Mg distances of BChl a within RC-LH1-PufX structure. Three groups of distances are observed, namely inter-, intra-LH1 BChl a distances, and 1-LH1/1-LH1’ Mg-Mg distance, the latter being the largest.

Its negatively charged headgroup participates in a localised electrostatic network, being in proximity to α-Trp45, α-Arg46, β-Arg49 as well as the neighbouring (N-1) β-Trp48 (**Fig. 4b**), apparently stabilising the headgroup, possibly serving as a Spno anchoring site. Overall, Spnos have a distinct and unique organisation within the LH1, either by shielding or exposing the BChl a molecules, working in concert to absorb distinct light and transfer the excitation energy absorbed. Other spheroidene subtypes are known to be present in *Rca. bogoriensis*^30^, but the achieved resolution of 2.9 Å does not allow further discrimination at the atomic level.

By applying C2 symmetry to reconstruct the cryo-EM map, each LH1 subunit is expected to exhibit conformational variation, as each monomer is treated as an asymmetric entity. Superposition of all 15 subunits highlights quite rigid structural conservation (higher in the centre), not only in terms of low α/β root-mean-square deviations, but also in terms of chlorophyll and carotenoid positioning (**Fig. 4b**). Despite this very low variation, we observe a gradually increasing trend in the LH1 as a function of structural variation of its subunits (**Fig. 4c**). The gradually increased dynamics observed may underlie subunit-specific conformational variation finely regulating the excitation energy path. Indeed, the conformational deviation from 1-LH1 as a function of subunit number **(Fig. 4c)** could be translated to the wobbling of the molecules in their resolved positions. The pattern roughly resembles the first harmonic vibrational pattern of a loosely confined (semi-free) vibrating bar. We postulate that this primary observation shows the energy transfer route between the ordered row of the light-harvesting complexes via a global resonance over the entire circumferential molecular framework. Such a mechanism may funnel the energy from either side of this framework to the RC. It is also intriguing that the 2-8 LH1 subunits are significantly less flexible as compared to the 9-15 LH1 subunits **(Fig. 4d)**. The LH1 subunit architecture is not only highly defined in terms of composition but also in terms of LH1-LH1 subunit proximity within the entire complex.

### Clustering of bacteriochlorophylls in LH1 reveals distinct excitation transfer pathways

Calculating Mg-Mg distances of LH1 chlorophylls across the RC-LH1 dimers discloses a bimodal distribution (**Fig. 4e**). Mg-Mg distances within an LH1 (α-BChl a-β-BChl a) are in the range of 9.0-9.3 Å, while those across LH1 subunits are substantially closer (8.3-8.8 Å) – similar distances were also reported for different number and composition of LH1 subunits in the *R. sphaeroides* RC-LH1 dimer grown in dark^23^ but no correlation similar to the one reported here was previously derived. The reported results may offer distinct vibrational frequencies at the quantum level to capture, emit or transfer quantas at different, discrete frequency or energy levels. Except for the 15-LH1, residing at the ubiquinone channel, also 7-LH1 exhibits high variation (**Fig. 4c**), which can be rationalized by its proximity in ultrastructural higher-order dimeric interfaces, visualized by freeze-fracturing at low resolution in endogenous membranes^9^ **(Fig. 1)**. Its position may allow cross-talk between neighbouring dimers. 7-LH1 also covers less interaction interface with the RC as compared to the other LH1 subunits, promoting a less bound state of an LH1 subunit. Overall, polypeptide conformational changes may further contribute to the captured, emitted, or transferred vibrational energy as previously shown for other photosystems with an exciton model based on quantum mechanics/molecular mechanics (QM/MM) simulations^31^. In the Mg-Mg proximity distribution (**Fig. 4e**), a point corresponding to the longest proximal Mg-Mg distance is revealed (d = 9.8 Å), mapping to BChls as that are each bound to the adjacent 1^st^ LH1 subunit from each of the RC-LH1 monomers, namely at the dimer interface. This distance is highly effective for electron transfer (d <10 Å)^32^, but also substantially larger than those across LH-1 subunits (∼0.5-1.5 Å longer), perhaps regulating excitation energy transfer within and across the monomers. Naturally, even slightly longer distances in the Å scale between adjacent pigments may confer substantially slower excitation energy transfer dynamics, therefore, regulating the migration path across the dimer. Possibly, this can be a species-specific variation that segregates quantum efficiency of energy trapping within each of the monomers, *i*.*e*., faster, and across the dimer, *i*.*e*., slower.

### PufX interacts with key proteins and pigments within the RC-LH1-PufX interface

Zooming into the structure of the 1-LH1/1-LH-1’ dimer, not only the proximal distance is larger than the rest **(Fig. 4e)**, but also residing orientations of resolved molecules are unique. One Spno is absent because it is replaced by interactions with PufX **(Fig. 5a)**. In addition, the phytol tail of the proximal BChl *a* undergoes a conformational change and ends up facing the periplasmic side of the RC-LH1 dimer (**Fig. 4b, 5a)**.

**Figure 5.**
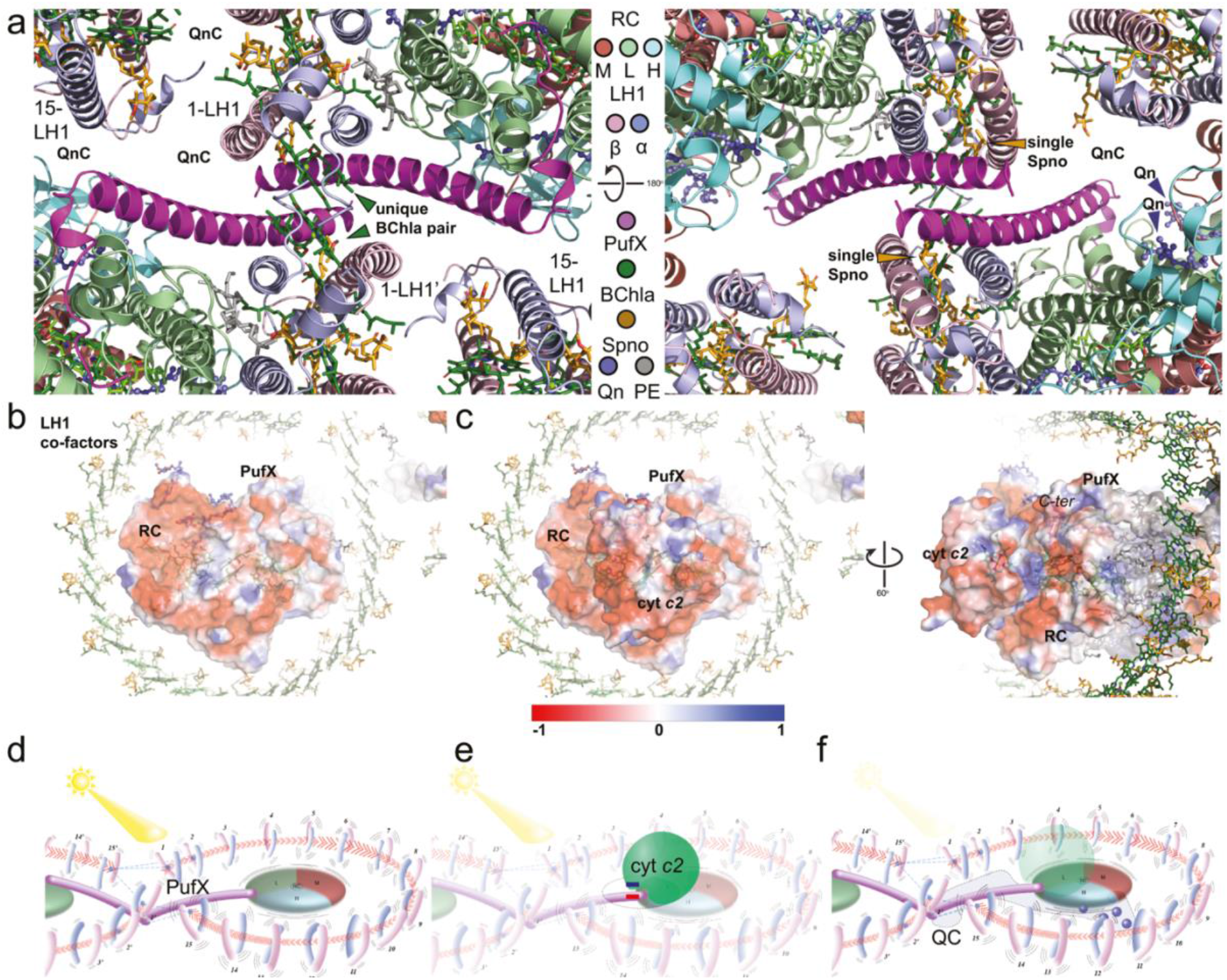
PufX localisation and contribution to RC-LH1-PufX stability and function. (a) PufX is dimerising and interacting within the dimer’s core interface; it also contributes to forming parts of the quinone channel (QnC) (b) Electrostatic surface potential map of the asymmetric unit of the RC-LH1-PufX. The *C-ter* of the PufX is shown to contribute to the highly negatively charged electrostatic surface potential. (c) When a cyt *c*_*2*_ is bound, electrostatic complementarity is visible, while PufX covers a substantial interface area, further contributing to the specificity and recognition of cyt *c*_*2*_ with the RC. (d-f) Schematics for the involvement of PufX in vibrational energy transfer (d), binding of electron donors/acceptors (e.g., cyt *c*_*2*_) (e) and structuring the quinone channel (f).

The *N-ter* region of PufX is localised in proximity, interacting with each LH1 subunit and, mostly, with their α-chains **(Fig. 5a)**. This confined interaction network among the 2 LH1 subunits and the PufX homodimer reveals and explains the previously hypothesised role for PufX in preventing certain components of the LH1 antenna from interacting with the RC in the QB pocket region, thereby blocking QB exit^33^.

In the reported RC-LH1 dimer, PufX shows an extensive, dense packing of interfaces with itself and the rest of the structure, involving Spnos, Bchl αs, lipids, antennas and two chains of the RC. This extremely dense packing of the dimer results in the formation of 7 interfaces (>100 Å^2^) per monomer with a total interface area of 4138,80 Å^2^ (**Extended Data Table 2**). PufX *C-ter* forms tight interactions with the RC (L subunit), covering 4.6% of its surface area. Various hydrophobic residues participate in the interactions forming a highly apolar, flat interface, absent hydrogen bonds (**Extended Data Table 2)**. Interestingly, its *C-ter* is exposed at the periplasmic face of the RC-LH1 dimer, proximal to the binding site of cytochrome *c2*, as well as various electron donors known from RC-LH1 monomers from other purple bacteria^34, 14^. PufX extends the surface area accessible to electron donor binding (**Fig. 5b**) substantially, further contributes to their attraction by exposing a highly charged *C-ter* tail (**Fig. 5b**), and when docked to cytochrome *c2* – a conformation known for *R. sphaeroides*^34^ – forms complementary electrostatics **(Fig. 5c)**. Such complementary electrostatics are known for protein-protein interactions with increased kon rates and contribute to interaction specificity^35^. Next, an overall stable interface of PufX *N-ter* with the 1^st^ LH-1 subunit is observed (1348,6 Å^2^) formed by two interfaces spanning both polypeptides forming the LH-1 subunit (890,4 Å^2^ and 458,2 Å^2^) buried by interacting with α and β chains, respectively (**Extended Data Fig. 4a)**. These interfaces have distinct physical-chemical properties.

Although apolar interactions majorly form both, a distinct hydrogen bond is observed (d=3.7 Å) between PufX (NH2 of Arg16) and the β chain of the 1^st^ LH-1 subunit (OG of Ser26). Overall, less polar residues are buried within the PufX-α chain interface than the one formed by PufX and the β chain. There are even other proximal protein chains to the PufX exhibiting hydrophobic interactions via their side chains and include (a) the Leu35-Val37 pair between PufX and the symmetrically-resolved 1^st^ LH-1 β chain (**Extended Data Fig. 4b)** and (b) the Met62-Pro305 pair (**Extended Data Fig. 4c)**, located at the termini of PufX and the M subunit of the RC, respectively. The back-folded PufX *C-ter* also displaces charged amino acids to accommodate the Met62 **(Extended Data Fig. 4c)**. The last protein interface is formed between the PufX monomers, covering 474,8 Å^2^, where the *N-ter* residues Asn17-Leu24 are involved, with the side-chains of Asn17, Met20 and Ser21 mostly contributing to the formed surface (**Extended Data Fig. 4d)**. The interactions mentioned above of PufX with RCs, LH1 and with itself clearly shows stabilisation of the RC-LH1 dimer via hydrophobic interactions residing in relatively flat, independent interfaces. These independent interfaces of distinct sizes regulate non-additive energetics to protein-protein interactions^36^, regulating affinity collectively. Apolar interactions regulate the stability of permanent complexes with very high affinities^35^.

Each PufX separately interacts with critical co-factors involved in the absorption and/or regulation of excitation energy: (a) the α-chain bound BChl, (b) its corresponding Spno, and (c) an L-chain phospholipid (3-*sn*-phosphatidylethanolamine, 8PE/3-sn-PE), covering interaction surfaces of 133 Å^2^, 326,2 Å^2^ and 162,4 Å^2^, respectively. The BChl tail (C4-C8) is stabilised via PufX residues Met23-Ala27, where hydrophobic stacking with the side-chain of Met23 is observed (**Extended Data Fig. 4e)**. Met23 and Leu24 are simultaneously involved in the dimerisation of PufX and the BChl tail stabilisation (**Extended Data Fig. 4e**), therefore, probably, having a dual role in assembling and supporting the RC-LH1 dimer. Next, the single Spno passing through the α/β chains of the 1^st^ LH1 subunit also mediates the interaction between PufX and the α chain of the antenna, bridging the components (**Extended Data Fig. 4e**). Again, the *N-ter* (Leu15-Met23) is involved in the interaction, where the Met23 side-chain of PufX is in direct proximity, facing the Spno. Therefore, this Spno is central, connecting the 2^nd^ LH1 to the 1^st^ LH1, the LH1-embedded BChls and Spno, and PufX (**Extended Data Fig. 4a, d-e)**. Interactions with the unsaturated tails of the 3-sn-PE are mediated by two consecutive Leu residues (Leu33 and Leu34), as well as a proximal Ile (Ile37). Interestingly, lipids significantly affect intra- and inter-subunit energy transfer efficiency and clustering^37^ with PE being highly advantageous for the self-assembly of LH2 into networks^34^. Finally, PufX contributes to the formation of the quinone channel **(Fig. 5a, Extended Data Fig. 4f)**.

All above-mentioned localised, packed networks of hydrophobic interactions are, therefore, responsible for the reorganisation of the BChl a conformations and bring the BChl a molecules bound to the 1^st^ LH1 antennas from each RC-LH1 complex in relative proximity (9.8 Å Mg-Mg distance), with possible implications in the differential efficiency of transfer within and across the RC-LH1 dimer. To conclude, we observed the involvement of PufX in the electrostatic attraction of electron donors/acceptors, e.g., cyt *c2* **(Fig. 5e)**; (b) structuring the quinone channel **(Fig. 5f)**.

### RC-LH1-PufX dimer

Overall, the resolved RC is composed of the L, H, and M protein subunits, protein U^38^ (also named protein Y^23^ whose sequence is different from that of *R. sphaeroides*) 4 BChl a, 3 Bacteriopheophytin a (BPhe a), 5 quinones (Q-10), 1 Spno, 4 PE lipids (3-sn-PE) and a Fe atom **(Fig. 6a, Extended Data Fig. 6)**. The resolved core RCs have a very conserved structure, highly similar to other described RCs from purple bacteria^7, 23, 38, 39^. Also, protein Y is resolved in the *Rca. bogoriensis* RC-LH1 dimer, establishing a parallel helix to the β chains of the 13^th^ and 14^th^ LH1 subunits, forming a trimeric helical bundle around the Spno charged tail (**Extended Data Fig. 7**). Its *C-ter* is proximal to a resolved Ubiquinone (U-10), the latter connecting the β chains of the 11^th^-13^th^ LH1 subunits with protein Y and extending the interaction network to the other known quinone molecules and the reaction centre. Notably, a bulky Trp residue of Protein Y is in proximity to one of the four BChl a tails resolved in the RC (**Extended Data Fig. 7**). A Spno molecule is resolved tightly embedded within the RC M chain, in interacting distance to the BChl a. Finally, an additional BPhe a is unambiguously resolved at the same conserved position as described earlier for the monomeric RC-LH1 from *Rba. veldkampii*^39^ ((**Fig. 6a**). This finding was never previously observed for any RC-LH1 dimer but given the origin of Rhodobaca (Ethiopia^26, 40^), a possible physiological role of this new pheophytin would be related to photoprotection, by rapidly quenching the higher excited states of the reaction **(Fig. 6a-c)**.

**Figure 6.**
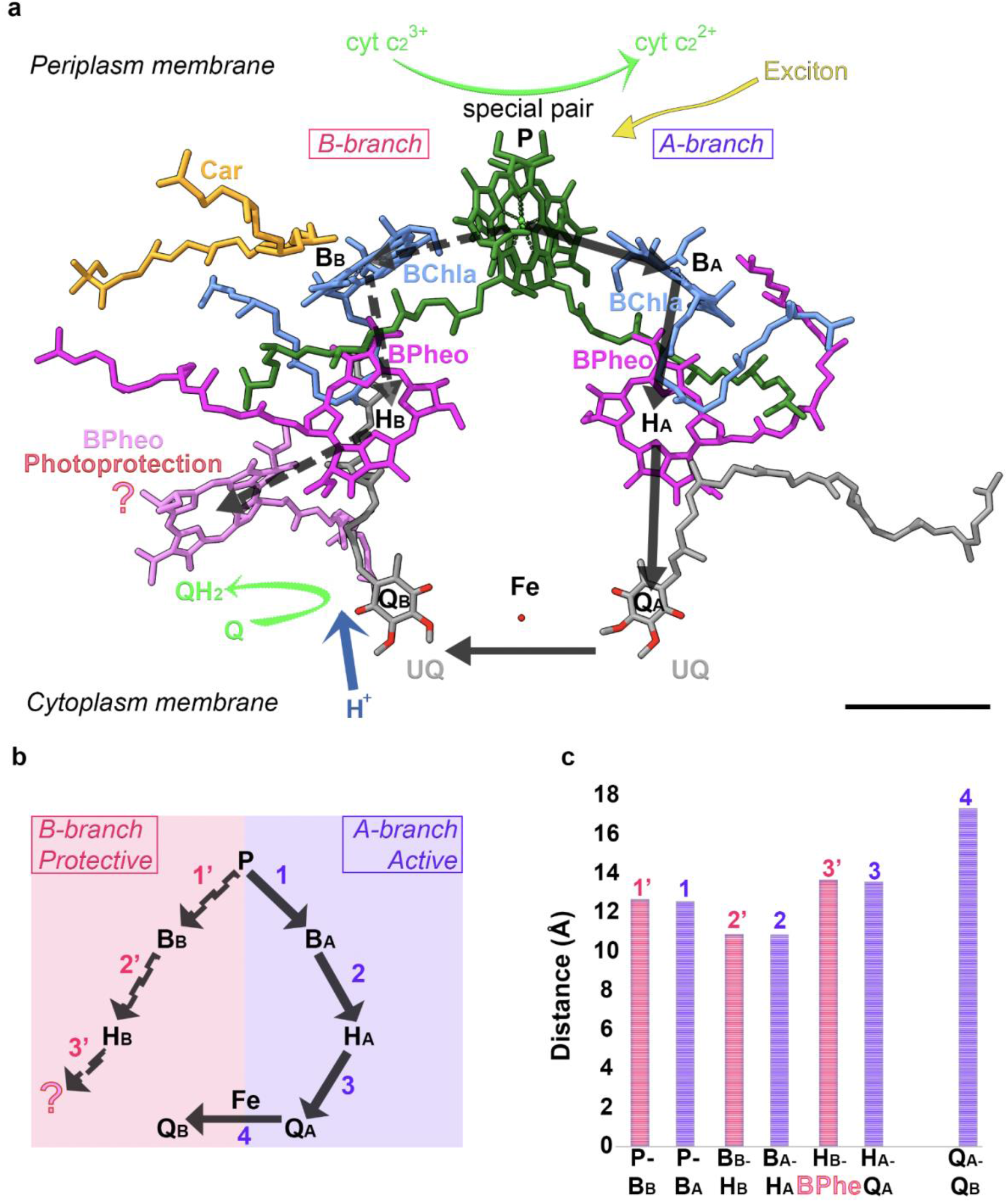
The organisation of the RC co-factors within the RC-LH1-PufX. (a) Visualisation of all co-factors and the RC of the RC-LH1-PufX. The scale bar = 1nm. (b) Pathways involved in electron transfer within the resolved RC. *A-branch (Active)* is coloured in pale mauve, *B-branch (Protective)* in pale pink; (c) Distances in angstrom between co-factors participating in *A-branch (Active)* and *B-branch*. The co-factor pairs are numbered according to panel (b). The bars’ colour indicating A/B branches are coloured correspondent to (b). Additional BPheo in pair HB-BPheo is coloured pink.

By clearly resolving all protein components and many novel co-factors for the *Rca. bogoriensis* RC-LH1 dimer, various electron transport paths are feasible within RC (**Fig. 6a-c**). The structure, therefore, proposes a unique mechanism for the capture and transfer of excitation energy within a wild-type, dimeric RC-LH1 photosynthetic apparatus. In the LH1, an unprecedented number of carotenoids (*n*=58) are asymmetrically bound and closely interact with the chlorophyll pairs (n=60). Their structure, architecture and proximity are highly conserved. Their collective excitation energy corroborates transfer in a highly efficient manner within a monomer. This is because within a monomer, this excitation energy, due to the closer proximity of BChl *a* molecules across and, not within LH1 subunits, has a higher probability to flow and arc around each monomer. The flow of excitation energy can also occur within the dimer, as evident by the close distance of the 1^st^ LH1 subunits for each monomer; However, this distance is larger than those identified within a monomer, and therefore, slightly lengthier energy transfer might occur. In addition, we cannot exclude an additional path that crosses the 1^st^ LH1 subunit of the RC-LH1 monomer and the 15^th^ subunit of its symmetric counterpart. Electron transfer in biologically relevant timescales may occur even at distances ∼2.5 nm^41^, which are well below the ones observed in the RC-LH1 structure between the corresponding BChl a Mg atoms. Ultimately, key energy transfer paths from the LH1 to the RC might be hypothesised based on three major relevant locations within the RC-LH1 dimer. One is via PufX, bridging to the RC and the residing Pheo a and BChl *a*, being also in vicinity to ubiquinones at the binding surface of electron acceptors.

## Conclusion and Perspectives

This study reveals the high-resolution structure of the RC-LH1-PufX dimer present in the bacterium *Rhodobaca bogoriensis* strain LBB1. Indeed, the structural data obtained during the last years on this complex came exclusively from the Rhodobacter family and, in particular, from *Rhodobacter sphaeroides*. It was, therefore, fascinating to compare the structures of these two bacteria. Our complex shows precisely the interaction of the two PufX polypeptides, whose role is essential to the supramolecular formation of the dimer. We find differences in the number of LH1 subunits, 15 in *Rca. bogoriensis* while *R. sphaeroides* has only 14. Moreover, an additional pheophytin is present along the B-branch of RC co-factors whose role remains to be elucidated. This branch is less active under normal growth conditions than the A-branch and could be solicited under specific conditions. One hypothesis considered for its function could be its involvement in photoprotection by serving as a sink for absorbing higher energy excited states by co-factors of the active branch reaction centre. Finally, a Y polypeptide is found between the LH1 antennae and the reaction centre. The presence of a tryptophan around the 9th amino acid is intriguing, considering the multiple functions of this large amino acid. This Y polypeptide could serve as a gateway for the quinones’ correct entry and/or exit from RC-LHCI or allow the quinones to accumulate in the right place.

*Open or closed ring?* The presence of PufX is essential for anaerobic photosynthetic growth in *R. sphaeroides*. The complete closure of the LH1 antenna around the RC in the PufX-deficient mutant inhibits quinone diffusion (the antenna closes poorly around the RC) and thus electron transfer between the RC and the *bc*1 complex^28^. This explains the inability of this mutant to grow under anaerobic photosynthetic conditions where the amount of oxidised quinones is meagre (1 quinone for about 50 quinol molecules), so they need to be re-oxidized rapidly, a prerequisite for efficient cyclic electron transfer.

Photosynthetic growth is restored in the PufX-mutant when the quinone pool is partially oxidised by the addition of an electron acceptor such as TMAO or DMSO^42^. These results imply that neither the supramolecular organization of the photosynthetic apparatus nor the open structure of LH1 in *R. sphaeroides* are necessary for cyclic electron transfer but suggest that the open cycle of LH1 appears to be necessary only when the quinone pool is fully reduced. Why is such supramolecular organisation (S-Shape) necessary in some species when the quinone pool is almost completely reduced? Because under such anaerobic growth conditions, the only oxidised quinone molecules are formed at the *bc*1 complex after charge separation. The concentration of oxidised quinones is shallow, and this situation implies that the quinones reduced by electron transfer must be oxidised very quickly. The opening of the LH1 antenna induced by PufX will facilitate the reduced quinones’ exit and recycle them more quickly as the dimeric centres have many more *bc*1 complexes (RC/*bc*1 ratio equal to 2) than the monomeric centres with closed LH1 antenna. This is observed with the dimeric reaction centre of *R. sphaeroides*^*4*^ and, possibly, *Rca. bogoriensis*. It is possible that for some species, the CRs with closed LH1, the physiological conditions in native membranes are such that, even under anaerobic conditions, the quinone pool is partially oxidised.

## Methods

### *Rca. bogoriensis* LBB1 culture and negative staining/freeze-fracturing EM

Wild-type *Rca. bogoriensis* strain LBB1 was isolated from soda lakes in the African Rift Valley by M T Madigan^26^ and kindly provided. Photoheterotrophic growth was carried out in anaerobiosis under continuous light (150 μE/m^2^/s) at 30°C in modified Hutner’s medium. For negative staining freeze-fracture, sample preparation, visualization and analysis were performed as in Semchonok et al^9^ without modifications.

### Isolation of the dimer followed the previous procedure^28^ with minor modifications

For the isolation of photosynthetic membranes, freshly harvested cells were collected by low-speed centrifugation at 8000 × *g*, washed with 50 mm Tris-HCl (pH 8) and 1 mm 4-(2-aminoethyl) bezenesulfonyl fluoride, and subjected to three cycles with French press at 16,000 p.s.i. The resulting unbroken cells and debris were removed by centrifugation at 20,000 × *g* for 30 min at 4 °C. The photosynthetic membranes were then pelleted at 200,000 × *g* for 90 min and resuspended in 50 mm Tris-HCl (pH 8) and 1 mm 4-(2-aminoethyl) bezenesulfonyl fluoride ^28^.

For the preparation of dimer complexes (see below), the membranes were treated with 3 M NaBr, followed by two washes with 50 mm Tris-HCl (pH 8) to remove membrane-attached proteins. In all experiments, the membranes were suspended in 50 mm Tris-HCl (pH 8.0). For the isolation and purification of photosynthetic complexes, the photosynthetic membranes were solubilized with 1% β-dodecyl maltoside for 15 min at 4 °C and then ultracentrifuged at 200,000 × g for 30 min to remove the insoluble material. The solubilised complexes were laid on a continuous density sucrose gradient generated by the freeze-thawing technique. The sample contained 50 mm Tris-HCl (pH 8), 0.5 m sucrose, and 0.03% β-dodecyl maltoside and ultracentrifuged at 200,000 × *g* for 15 h at 4 °C. Three pigment-containing bands were thus separated and ascribed (from top to bottom) to monomeric and dimeric RC-LH1 complexes. The latter fractions were carefully removed using a syringe and were applied to a Mono Q anion exchange fast protein liquid chromatography column (Amersham Biosciences) pre-equilibrated with 50 mm Tris-HCl (pH 8), and 0.03% β-dodecyl maltoside. After washing with 5 ml of buffer, the complexes were eluted with a 30-ml linear gradient of 0–500 mM NaCl at a 0.5 ml/min rate. HPTLC coupled to ESI-Tandem MS was used for identifying phospholipids associated with the dimer^43^.

### Sample preparation for electron microscopy

The sample was shipped in dry ice. Upon arrival, freeze-thawing was performed, and Bradford concentration was measured for different dilutions. The final concentration was 3.1 mg.ml^-1,^ which was chosen for direct cryo-EM grid preparations. Immediately before sample preparation, the sample support grids were glow discharged in low-pressure air with 15 mA current in a PELCO easiGlow. Cryo-EM grids were prepared by plunge-freezing in liquid ethane on a Vitrobot Mark IV (Thermo Fisher Scientific, USA). The glow discharge and plunge-freeze parameters are listed in **Extended Data Table 1**.

### Electron microscopy

The dataset was collected on a Glacios (Thermo Fisher Scientific, USA) 200 kV electron microscope equipped with Falcon 3EC direct electron detector (Thermo Fisher Scientific, USA). Movies were acquired with EPU 2.11 software, one image per hole and saved as gain-normalized MRC files. The acquisition parameters are listed in **Extended Data Table 1**.

### Image processing

The processing workflow trees for the investigated subsets are presented in **Extended Data Fig. 1**. The data processing was performed in Scipion 3.0 framework^44^. Dataset was motion-corrected with MotionCor2 v1.4.3^45^ using 5 × 5 patches, no frame grouping, B-factor 500, with saving of dose weighted and averages. The CTFs were fitted on the dose-weighted averages using Gctf v1.06^46^ with a 20–3.5 Å resolution range and a 1024-pixel box. The particle picking was performed by xmipp3^47^ – manual//automated particle picking and SPHIRE-CRYOLO^48^ picking using for the training the picked coordinates from xmipp3-manual picking protocol completed previously. The resulted coordinates were then compared, the repeated positions were removed with xmipp3 – picking consensus protocol for the radius value of 50 pixels. The particles were extracted with a 384×384 box size in pixels. The 2D classification was done with cryoSPARC (plugin v.3.0)^49^ – 2D classification protocol (Scipion 3.0 plugin v.3.0). Particles from the best-resolved 2D classes were subjected to RELION 3D initial model protocol^50^ with C2 symmetry and default parameters.

Then the initial particle sets (0.96 A/pix, box 384) were applied for RELION 3D classification with default regularisation parameter (tau fudge) values of T = 4 for 3D classifications. A 20 Å low pass filtered internal reference and “fast subsets” (small random particle subsets in the initial iterations) were used to prevent bias propagation and to provide featureless references for false positives and contaminants to be classified into. Particles from the best-resolved class of the initial 3D classification were subjected to cryoSPARC (plugin v.3.0) 3D non-uniform alignment^51^ (plugin v 3.0) with C2 applied symmetry that included defocus refinement with +/-4000 defocus search rate, global CTF refinement (including enabling of Fit Tilt, Fit Trefoil, Fit Spherical Aberration, Fit Tetrafoil parameters). The refined 3D map was subjected to RELION – post-processing step using the mask generated during the previous refinement step in cryoSPARC protocol (plugin v.3.0).

The 30 e^**-**^/Å^2^ exposure frame subset of RC-LH1 used the first 30 of 120 movie frames in the motion correction and Bayesian polishing steps. Then the cryoSPARC 3D non-uniform refinement (plugin v.3.0) with C2 and C1 applied symmetry was executed with the reference and mask obtained during the previous refinement step in cryoSPARC (plugin v.3.0). The outcome resulting dataset was subjected to cryoSPARC local resolution (v3.2). Local resolution was estimated in cryoSPARC (plugin v.3.0) using a windowed FSC method^52^ with FSC threshold 0.5 and with step size 1 Å; cap local resolution estimation and FSC weighting parameters were enabled and, finally, the map was visualised in ChimeraX^53^. Map sharpening was done in cryoSPARC (v3.2) by applying the overall B-factor estimated from Guinier plots. By default, masks were generated during 3D non-uniform alignment in cryoSPARC (v3.2). The visualisation was performed with Chimera^54^, ChimeraX^53^ and PyMOL^55^.

### Model building

Iterative cycles of building and refinement on the working model were done using Coot ^56^ and Phenix’s “real space refine” tool^57^, respectively, starting from a homologous RC-LH1-PufX monomer^39^. Ligands already present in Protein Data Bank^58^ were added and placed manually in Coot using the following PDB ligand ids: 8PE, BCL, BPH, PEF, U10, UQ5, and UQ8. The novel carotenoid Spheroidenone (PubChem^59^ CID: 5366412) was built in the canonical SMILES string in AceDRG ligand builder^60^ from the CCP4 suite^61^. Ligand restraint and libraries, used for real-space refinement, were generated using the Phenix eLBOW tool^62^.

### Protein Y identification

We determined the protein candidate based on the electron density map to include a tryptophan residue between positions 5 and 20 in the amino acid sequence and a display size between 50 and 100 amino acids. As no protein-coding sequence (CDS) from the Puf operon could match, we thus extend our candidate research on the complete genome of *Rca. bogoriensis* (Bioproject: PRJNA224116, Assembly GCF_014197665.1). A program was written in R language using Rstudio, which retrieved a specific protein sequence by restraining the size of the candidate and selecting the position of a specific residue. The Biological Sequences Retrieval and Analysis (seqinr) library was used^63^ and the sequence of the Protein Y (Protein U) identified based on the bioinformatic analysis and the cryo-EM map was: >NZ_JACIGD010000002.1_prot_WP_071480931.1_1389[locus_tag=GGD87_RS07010] MEHTVVSHSWLIETLGEVRKYAARNGLPALAEHLEQAIHLAHIEQATLEGQDSEGKKRSNGENGPDDQSR.

## Supporting information

SuppFigs_Tables

## Acknowledgements

This work was supported by the Federal Ministry for Education and Research (BMBF, ZIK program) (Grant nos. 03Z22HN23, 03Z22HI2 and 03COV04 to P.L.K.), the European Regional Development Funds for Saxony-Anhalt (grant no. EFRE: ZS/2016/04/78115 to P.L.K.), funding by Deutsche Forschungsgemeinschaft (DFG) (project number 391498659 and RTG 2467), and the Martin-Luther University of Halle-Wittenberg. We thank all members of the Kastritis Laboratory for valuable discussions. The authors wish to acknowledge Dr Thomas Hoffmann of Thermo Fisher Scientific for his valuable suggestions on cryo-EM data acquisition. The authors thank Sacha Peschoux and his programming skills that contributed to protein Y identification; MT Madigan and his team kindly provided the strain, and Jerome Lavergne for contributing to the advancement of knowledge about *Rca. bogoriensis*.

## Author contributions

C.J., D.A.S. and P.L.K. conceptualised and initiated the project. C.J. and Q.C. performed protein purification. F.L.K. and D.A.S. performed grid preparation and vitrification. F.H. set up and supervised cryo-EM data collection. F.H. and D.A.S. did the data collection. D.A.S. performed image processing. M.I.S., Y.S. and C.T. performed the model building. F.H, F.K and P.L.K assisted with cryo-EM data collection and advised in sample and data collection optimisation. P.L.K. and C.J. wrote the manuscript with the input of all other authors.

## Data availability

The 3D maps will be available at the EMDB database (accession code EMD-XXXXX and EMD-YYYYY) and the molecular model of the RC-LH1-PufX dimer at the PDB database (accession code PDB ID: XXXX)). All accessions will be freely available for downloading after revision.

## Code availability

All unpublished code and scripts used in this study are available upon request.

## Ethics declarations

### Reporting Summary

Further information on research design will be available in the Nature Research Reporting Summary submitted with this draft.

