## Supplementary material for "Cryo-EM structure of the *Rhodobaca bogoriensis* RC-LH1-PufX dimeric complex at 2.9 Å": SuppFigs_Tables

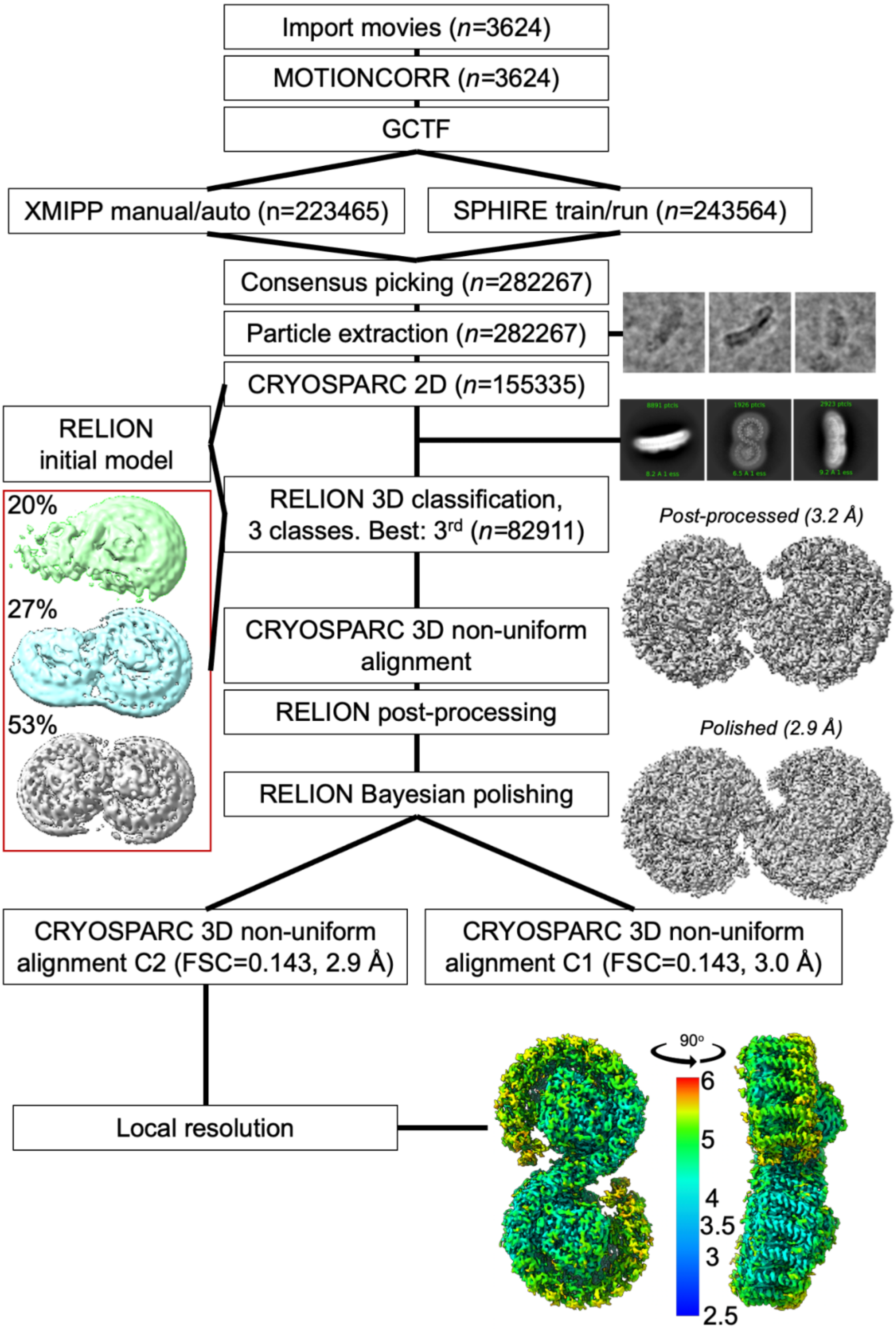

**Extended Data Fig. 1. Workflow for the cryo-EM map calculation was performed utilizing the Scipion 3.0 image processing framework, including RELION 3.1 and cryoSPARC 3.0.**

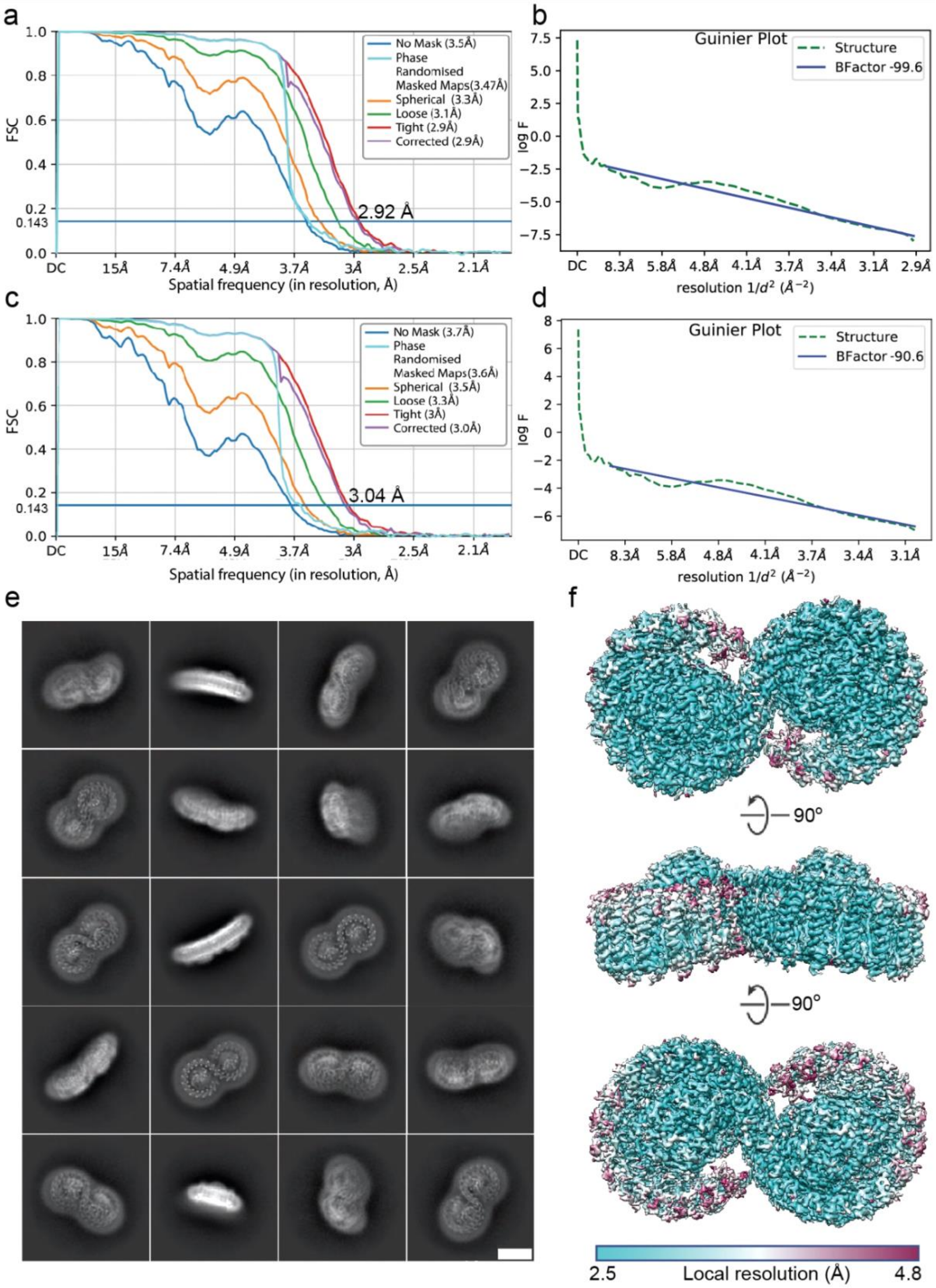

**Extended Data Fig. 2. Validation of cryo-EM data**. (a) Gold-standard Fourier Shell Correlation FSC of the final C2 symmetric reconstruction; (b) Guinier plot for the final C2 symmetric reconstruction; (c) The same as (a) but for the final C1 reconstruction; (d) The same as (b) but for the final C1 reconstruction; (e) Best class averages from cryoSPARC 3.0 2D classification. Various views are visible. Scale bar is 10 nm; (f) Local resolution estimation showing a variation from 2.5 Å to 4.8 Å, the lowest mostly localized at the late antennas and the dimeric interface of the complex. (f) from the top — periplasmic, in the middle - parallel to the membrane plane and cytoplasmic (at the bottom) views.

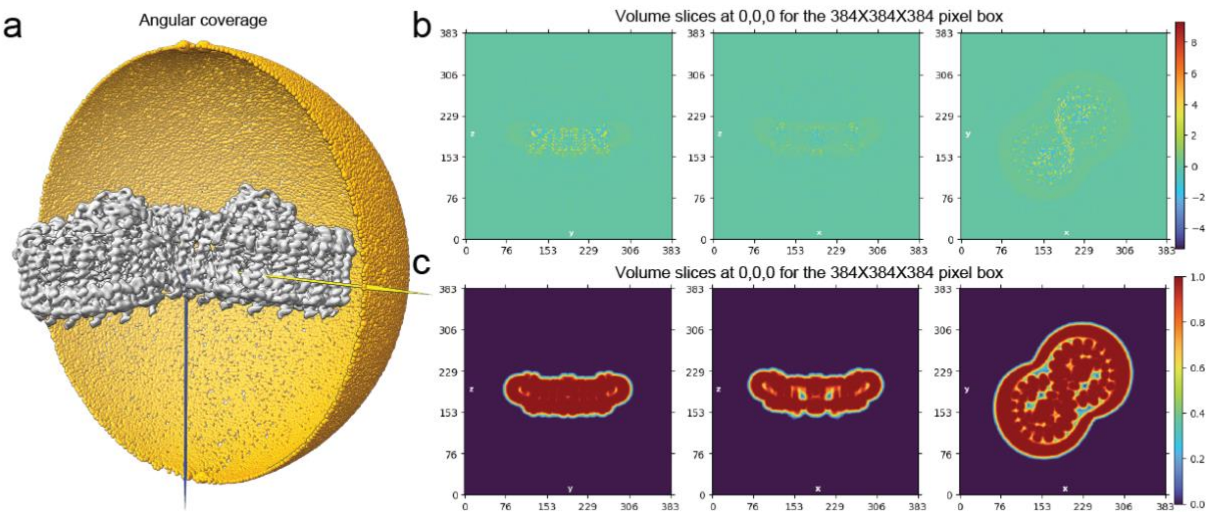

**Extended Data Fig. 3. Extended map validation and statistics.** (a) Angular distribution of final C2 map; The yellow half-spheres and their size represents the angular distribution of reconstructed particles in the C2 map of the RC-LH1-PufX complex. (b) Projections at the center of origin for the C2 map of the RC-LH1-PufX complex. (c) Projections at the centre of origin for the mask applied to calculate resolution for the RC-LH1-PufX complex.

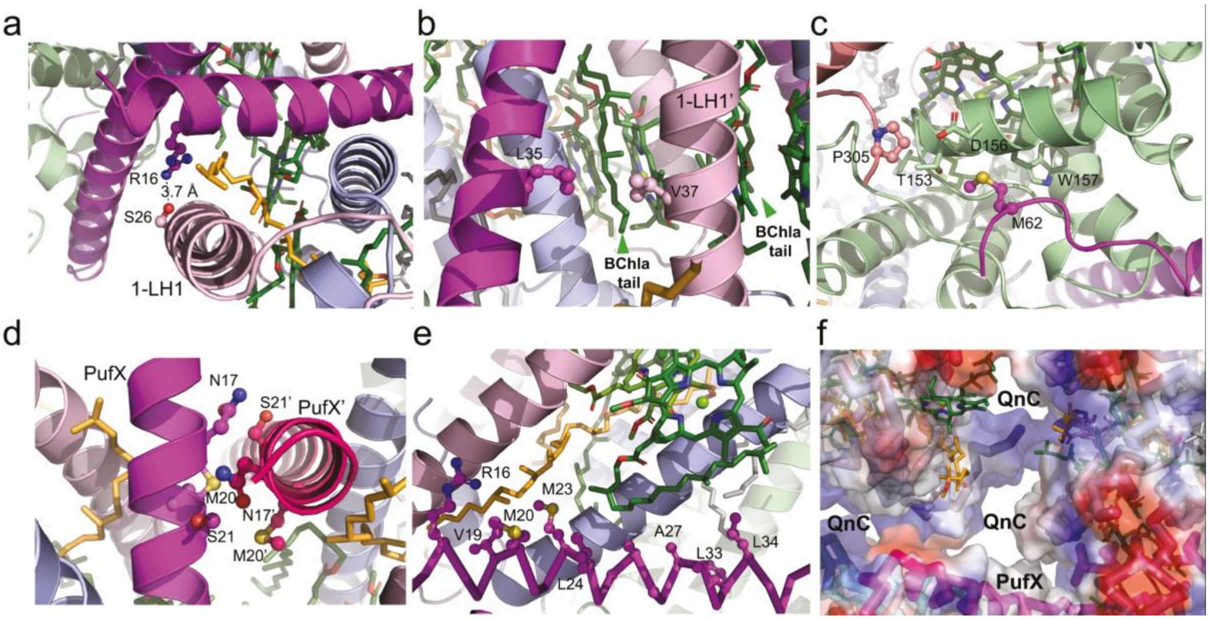

**Extended Data Fig. 4. Interactions of PufX within the RC-LH1-PufX cryo-EM model.** (a-e) Interactions of PufX with surrounding amino acid residues and pigments; For details, see main text. (f) PufX contributes surface area in the quinone channel (QnC).

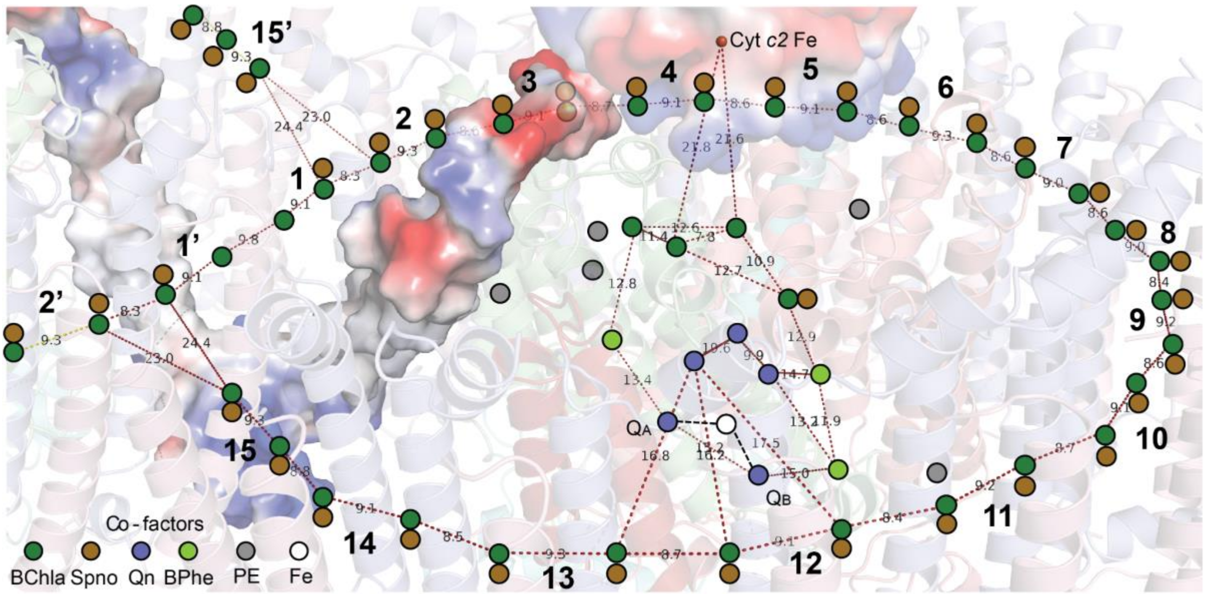

**Extended Data Fig. 5. Placement of approximate centroids for co-factors identified within the RC-LH1-PufX complex along with distance calculations across co-factors.** PufX is shown with a calculated electrostatic potential map, the superimposed cytochrome $c_2$ with a more transparent surface, and the iron of its heme is also shown in atom representation. The colour code for co-factors is described in the insert.

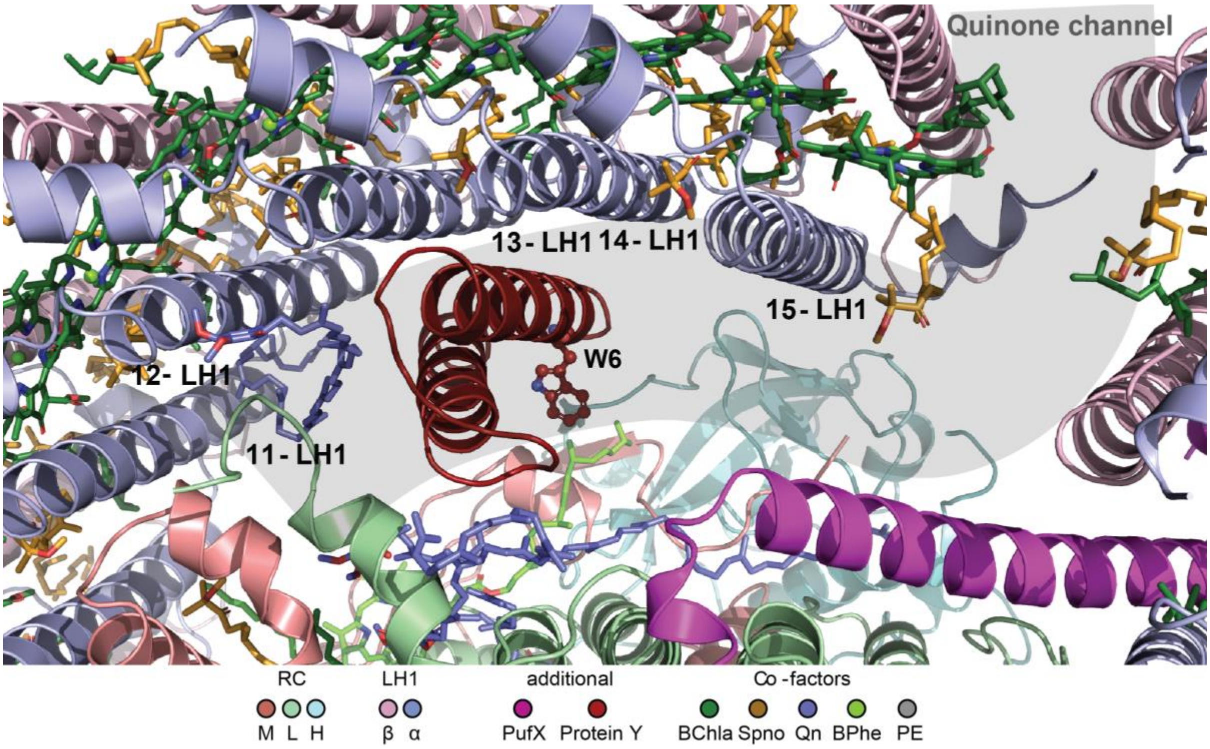

**Extended Data Fig. 6. Protein Y and its localization between the RC and the LH1. View from the periplasmic side of the membrane.** For details, see text.

**Extended Data Table 1. Sample preparation, Microscopy, Data processing and validation.**

| | #1 RC-LH1-PufX (C2) (EMDB-xxxx) (PDB xxxx) | #2 RC-LH1-PufX (C1) (EMDB-xxxx) |
|---|---|---|
| **Sample preparation** | | |
| Concentration [mg/ml] | 3.1 | |
| Sample volume [µl] | 3.5 | |
| Grid type | Quantifoil R 2/1 Cu 200 | |
| Glow discharge time [s] | 25 | |
| Glow discharge current [mA] | 15 | |
| Glow discharge sample polarity | Negative | |
| Glow discharge atmosphere | residual air | |
| Glow discharge pressure (Pa) | 40 | |
| Blotting chamber temperature [°C] | 4 | |
| Blotting chamber humidity [%] | 95 | |
| Blot time [s] | 6 | |
| **Microscopy** | | |
| Magnification | 150 000 | 150 000 |
| Voltage (kV) | 200 | 200 |
| Focal length (mm) | 3.4 | 3.4 |
| Cs (mm) | 2.7 | 2.7 |
| Objective Aperture (µm) | 100 | 100 |
| Number of movies | 3624 | 3624 |
| Electron exposure (e–/Å$^2$) | 120 frames (1 e-/frame) | 120 frames (1 e-/frame) |
| Defocus range (µm) | -0.5 to -1.5 | -0.5 to -1.5 |
| Pixel size (Å) | 0.9612 | 0.9612 |
| Symmetry imposed | C2 | C1 |
| Initial particle images (no.) | 282267 | 282267 |
| Final particle images (no.) | 82911 | 82911 |
| Map resolution (Å)/ B-factor (Å$^2$)<br>FSC threshold | 2.9 / -99.6<br>0.143 | 3.0 / -90.6<br>0.143 |
| Map resolution range (Å) | n/a | n/a |
| **Data processing** | | |
| Refinement | | |
| Initial model used (PDB code) | 7DDQ | |
| Model resolution (Å)<br>FSC threshold | 2.9<br>0.143 | |
| Model resolution range (Å) | 2.9-3.2 | |
| Model composition<br>Non-hydrogen atoms<br>Protein residues<br>Ligands | <br>22587<br>2327<br>BCL: 34, LIG: 30, BPH: 3, 8PE: 3, UQ8: 2, FE: 1, U10: 2, PEF: 1, UQ5: 1 | |

| | |
|---|---|
| *B* factors (Å$^2$, min/max/mean)<br>Protein<br>Ligand | <br>0.21/170.18/54.09<br>0.11/161.24/50.22 |
| R.m.s. deviations<br>Bond lengths (Å)<br>Bond angles (°) | <br>0.008<br>1.508 |
| Validation<br>MolProbity score<br>Clashscore<br>Poor rotamers (%) | <br>2.35<br>14.32<br>3.13 |
| Ramachandran plot<br>Favoured (%)<br>Allowed (%)<br>Disallowed (%) | <br>95.57<br>4.43<br>0.00 |

**Extended Data Table 2. PufX interfaces within the RC-LH1-PufX dimer.**

| *Interface with:* | $N_{atoms}$ | $N_{res}$ | *Surface, Å*$^2$ | *Chain (PDB)* | $N_{atoms}$ | $N_{res}$ | *Chain (PDB)* | *Interface, Å*$^2$ | $N_{Hb}$ |
|---|---|---|---|---|---|---|---|---|---|
| RC, L subunit | 73 | 18 | 5353 | L | 95 | 30 | 16663 | 1629.2 | 3 |
| α-LH1 | 45 | 12 | /=/ | a | 48 | 12 | 4357 | 890.4 | 0 |
| PufX | 23 | 6 | /=/ | Y | 23 | 6 | 5354 | 474.8 | 0 |
| β-LH1 | 21 | 7 | /=/ | A | 24 | 8 | 4155 | 458.2 | 1 |
| Spno | 25 | 6 | /=/ | [LIG]a:102 | 15 | 1 | 1085 | 326.2 | 0 |
| Phospholipid | 5 | 3 | /=/ | [8PE]L:306 | 4 | 1 | 1034 | 162.4 | 0 |
| BChla | 8 | 3 | /=/ | [BCL]a:101 | 5 | 1 | 1093 | 133.0 | 0 |
| β-LH1' | 2 | 1 | /=/ | O | 2 | 1 | 4151 | 61.0 | 0 |
| RC, M subunit | 1 | 1 | /=/ | M | 1 | 1 | 18485 | 0.6 | 0 |
